# Identification of Novel Open Reading Frames on the Rabies Virus Antigenome Strand and Assessing Their Expression

**DOI:** 10.64898/2025.12.26.696564

**Authors:** Ankeet Kumar, Utpal Tatu

## Abstract

Rabies disease can be caused by several lyssaviruses, which are a group of bullet-shaped, enveloped viruses with single-stranded, negative-sense RNA as their genome. Lyssaviruses are known to produce a full complement of genomic RNA (cRNA or antigenome) during their lifecycle. The rabies virus (RABV) is the prototype of the genus and has a genome of 12 kilobases. The RABV genome is known to encode five proteins from the genomic strand. In this study, we examined the coding potential of RABV and identified 17 novel ORFs (nORFs): one on the genomic strand and sixteen on the antigenomic strand. Synteny analysis revealed that several of these nORFs were conserved across the RABV clades. Additionally, five of the nORFs possessed Kozak sequences, and trRosetta predicted well-folded three-dimensional domains in these nORFs. Using LC-MS data of rabies-infected canine brain samples, we identified high-confidence peptides corresponding to the two nORF-coded proteins, nORF9 and nORF12. Furthermore, the structural and sequence similarity analyses indicated that these domains share homology with proteins involved in signalling and transcription.

## Introduction

Rabies virus (RABV) belongs to the genus *Lyssavirus*, which is a group of negative-sense RNA viruses under the order Mononegavirales [1]. Several of these viruses are known to infect humans and cause a disease called rabies [2]. The disease spreads via the bites from an infected animal to other animals. It is 100% fatal if not treated in time [1]. The only treatment available is vaccination or a combination of vaccination and rabies immunoglobulins, based on the type of injury [3]. The disease has been known since ancient times, but it is still a huge issue for developing countries, and more than one-third of the human deaths reported around the globe are known to occur in India [4]. There are mainly two reservoirs for it, bats and dogs; dog-associated cases account for more than 95 % of the cases [4].

RABV genome is around 12 kb long, negative-sense RNA, and is known to harbour 5 genes in the order of 3’ Nucleoprotein (N), Phosphoprotein (P), Matrix protein (M), Glycoprotein (G) and Large (L) protein 5’, and these genes are transcribed into individual mRNAs by RNA-dependent RNA polymerase coded by L gene [5,6]. The L gene is the largest gene (6428 bp) in its genome, accounting for approximately 50% of the genome’s length. The G gene codes for a protein that is 524 amino acids long and is the host receptor-binding protein. Phosphoprotein is a multifunctional protein, which is annotated as a 297 amino acid-long protein. However, there is evidence that this protein also codes for four additional isoforms (P2-P5) [7], in addition to the canonical P protein (P1). One of the isoforms, called the P3 isoform, harbours a nuclear export signal (NES) and retains a nuclear localisation signal (NLS) [8,9]. This isoform is deemed to be very crucial for interfering with the expression of immune genes, as it shuttles between the nucleus and cytoplasm and can bind to the STAT protein [8].

However, whether other proteins express any isoform or if there are other unannotated ORFs present in the RABV genome is not understood hitherto. A complete antigenome is present during the rabies life cycle, and a study from 1998 shows that the RABV genome has the capability to express genes inserted into the antigenome [10]. In this study, we investigated the presence of novel open reading frames (nORFs) on the antigenome of RABV. We identified the presence of an additional 17 nORFs in the RABV genome, spanning both the canonical and antigenome, which were previously unreported. We used in-silico predictions and backed up this observation with proteomics data to demonstrate the expression and presence of peptides for nORFs. We also find that several of these nORFs are conserved across the RABV clades. Additionally, we employed multiple structural and sequence homology tools to investigate the function of these novel ORFs and gain insights into their potential roles in viral biology.

## Methods

### Finding all open reading frames in the reference genome of RABV

We used the ORF finder (https://www.ncbi.nlm.nih.gov/orffinder) [11] to assess the coding potential of RABV using its reference genome sequence (NC_001542). We used a cutoff of 150 bp and kept other options as the default in the ORF finder, which revealed the presence of twenty-two ORFs in the reference genome sequence.

### Conservation of novel open reading frames of RABV

We performed neighbour-joining phylogenetic analysis for the eight clades of RABV using RABV-GLUE (https://rabv-glue.cvr.gla.ac.uk) [12] clade-reference sequences. Furthermore, we utilised ORFfinder to predict the coding potential of these sequences, followed by identifying similar nORFs in Microsoft Excel and determining protein sequence similarity using EMBOSS-NEEDLE (https://www.ebi.ac.uk/jdispatcher/psa/emboss_needle) to understand the conservation of nORFs. We also used SimpleSynteny (https://www.dveltri.com/simplesynteny/) [13] to decipher the relation across these nORFs. The nORF sequences and their conservation are available as **Supplementary Table 1**.

### Identification of peptides corresponding to the novel open reading frames in the proteomics data

We identified signals for translation using the ATGpr (https://atgpr.dbcls.jp/) tool [14], which identified signals for protein translation for the nORFs; only 5 nORFs showed the signals. The file used for ATGpr is presented in **Supplementary File 1**. To understand if the nORFs coded proteins are translated during the infected condition, we used proteomics data from our lab [15] generated from infected canine brain isolates and analysed using Proteome Discoverer (v2.5) [16] to identify peptides corresponding to the nORFs. The database used for proteomics searches is available as **Supplementary File 2**.

### Understanding the role of the novel open reading frames

We employed structural homology search and sequential domain predictions to elucidate the potential role of these nORFs. We used trRosetta [17] to predict the structures, HHpred [18] to identify domains for the novel peptides, DeepFold (https://zhanggroup.org/DeepFold/) [19], Grand Average of Hydropathy (GRAVY) [20,21], HMI-PRED (https://interactome.ku.edu.tr/hmi/) [22] to identify similar domains or folds in other proteins, and also to predict the probable interactome of these proteins. The structures were visualised in PyMOL [23].

## Results

### RABV antigenome harbours open reading frames coding for small proteins

To investigate the coding potential of RABV, we used the ORF finder to identify the coding regions for RABV. In addition to the five canonical genes N, P, M, G and L, which are present on the negative-sense strand of the virus, the virus harbours other open reading frames. In total, an additional 17 nORFs can code for polypeptides of more than 50 amino acids. **Figure 1A** shows that most of these nORFs are present on the antigenome (reverse-strand) genome. Only one nORF (ORF3), coding a protein of 102 amino acids, is present in the canonical strand, while all other nORFs are present on the antigenome. **Figure 1B**, the median length of the nORFs is 77.94 amino acids, with the largest one coding for 130 amino acids and the smallest one coding for 51 amino acids.

**Figure 1.**
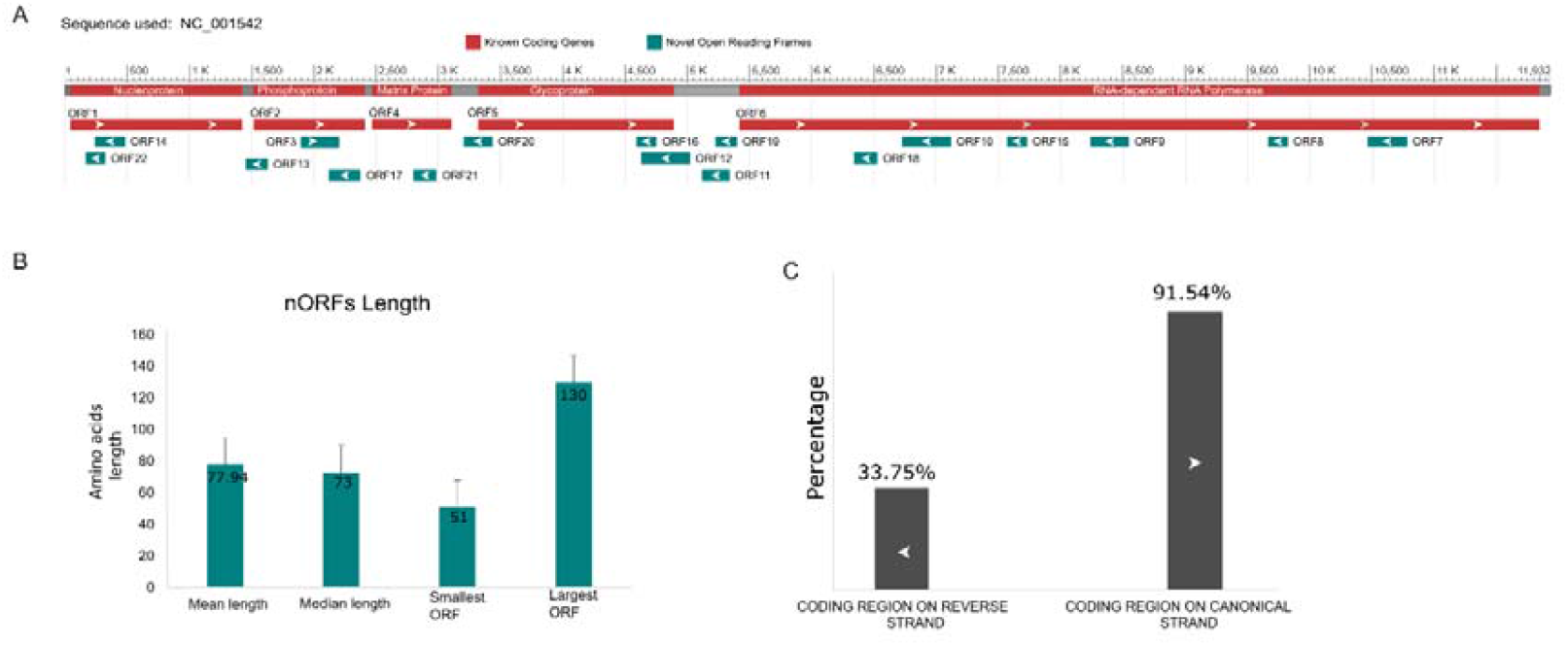
The coding potential of the rabies virus (RABV). A) The prediction of genes on the reference sequence (NC_001542) of RABV using a criterion of 150 bp as the smallest ORF. The arrow inside the ORF shows the direction of the coding region in reference to the annotated genome. B) The bar chart showing the length of nORFs with error bars. C) The percentage of the coding region on the canonical and the antigenome strand.

**Figure 1C** shows the coding region percentage on the canonical strand versus the antigenome. On the canonical coding strand, 91% of the nucleotides are involved in the coding region, whereas on the reverse strand, only around 34% of the region harbours coding potential for RABV (based on NC_001542). Fasta sequences of all the nORFs are presented in the **Supplementary File 3**.

### Several novel open reading frames are conserved across the clades of RABV

The conservation of the coding regions across the evolutionary scales is often associated with their functional importance [24]. We examined the evolutionary conservation of nORFs across the eight clades of RABV, which have diverged over a significant evolutionary timescale [25]. **Figure 2A** presents the neighbour-joining tree for the representative sequences, showing the relationship among the RABV clades. RABV phylogeny reveals two main branches, one leading to bat-related clades (Bats and RAC-SK) and the other to canine-related clades (Africa-2, Africa-3, Arctic, Asian, Cosmopolitan, and Indian Subcontinent).

**Figure 2.**
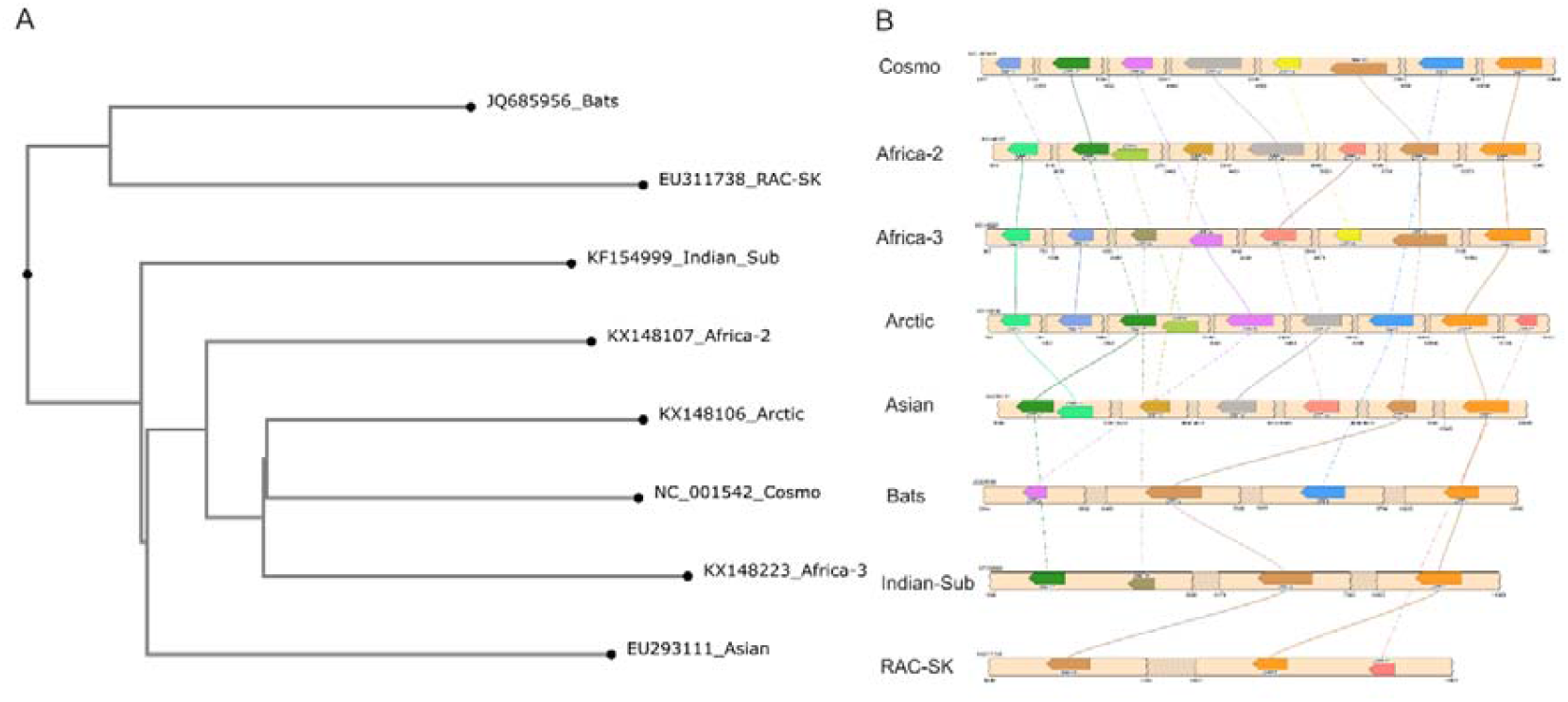
Neighbour-joining phylogeny and nORF similarity of RABV clades. A) A maximum-likelihood phylogeny of the eight lineages of RABV. B) A synteny map showing the conservation of several ORFs between and across the clades of RABV.

**Figure 2B** presents the conservation of several ORFs across the eight clades of RABV. The synteny map shows that ORF7 is conserved across all the clades. ORF10 is absent only in the Arctic clade, and ORF21 is only present in RAC-SK and Arctic. Whereas others differ in their presence across clades and are shared only among some lineages.

### Peptides corresponding to two nORF-coded proteins were identified in proteomics data

Out of the five candidate non-coding open reading frames (nORFs) that contained Kozak consensus sequences potentially indicating translation competence, **Supplementary Table 2**. We identified 6 peptides corresponding to two proteins, observed in the MS/MS (tandem mass spectrometry) spectra analysis. For nORF9, we observed four unique peptides (NSEFSTLSTSSYKAFLIASLSR, KDELNSSLR, LSLWVLDNIASR, and MIDSGIPKK), and for nORF12, we only observed two unique peptides (TPQGEVIFRTWIVER and RTPQGEVIFRTWIVER) presented in **Figure 3**. This provides direct proteomic evidence demonstrating that the predicted open reading frames are translated. **Table 1** shows the peptides found in the proteomics samples.

**Table 1.**
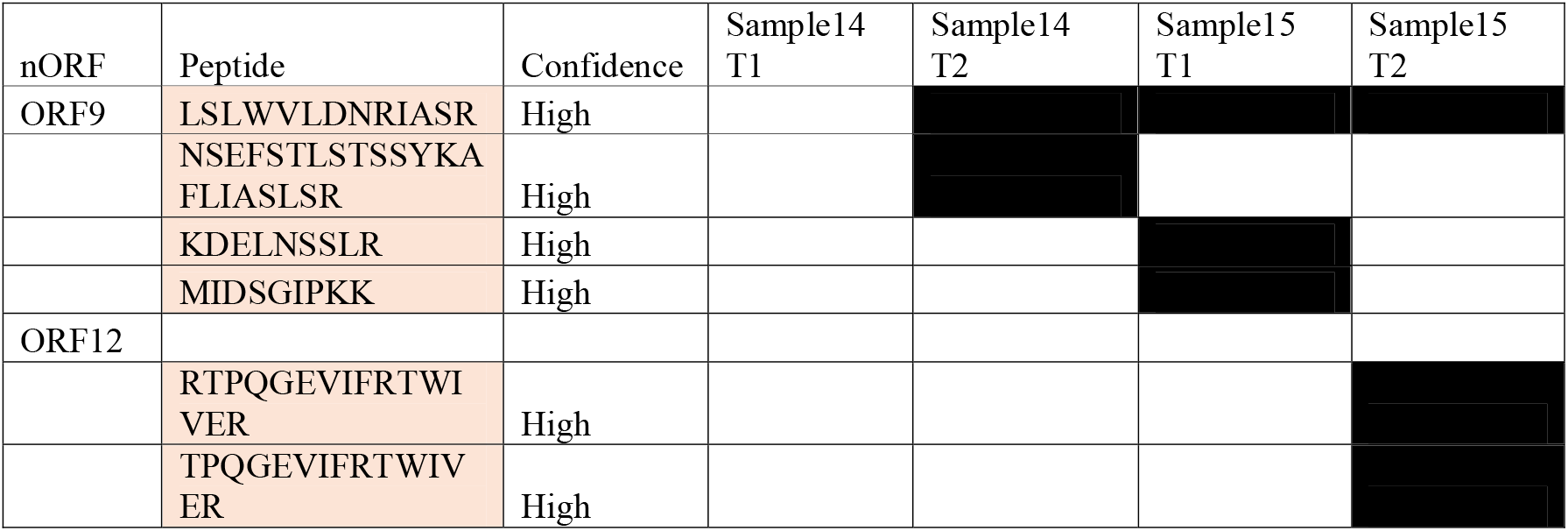
Peptides for nORFs in the proteomics of infected canine brain samples, with a cutoff of more than 2 unique peptides for the nORF.

**Figure 3.**
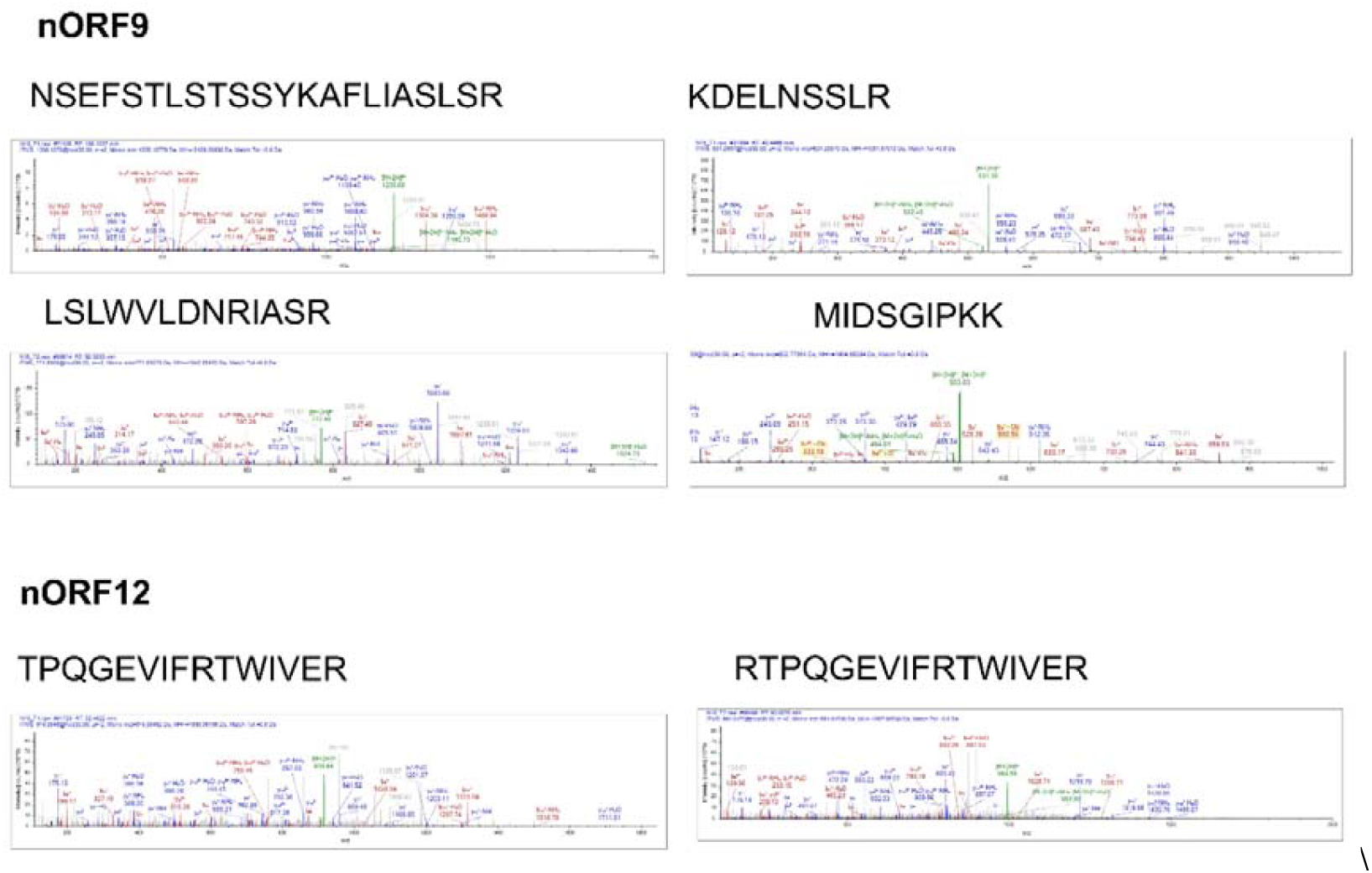
Peptide spectra verifying the translation of novel open reading frames (nORF9 and nORF12). Shown here are representative MS/MS spectra of the peptide sequences that exclusively mapped to nORF9 (NSEFSTLSTSSYKAFLIASLSR, KDELNSSLR, LSLWVLDNIASR, MIDSGIPKK) and nORF12 (TPQGEVIFRTWIVER, RTPQGEVIFRTWIVER). Annotations of the fragment ion series (b- and y-ions) confirm assignments of peptide sequences

### Structural homology and sequential homology-based functional assessment of the nORFs

The structures generated using trRosetta were used as an input for Foldseek to ascertain the functional domains using structural homology. **Figure 4** presents the structures of the 5 ORFs that showed translation signals. All of the novel proteins exhibit well-defined three-dimensional domains, whereas some parts of the proteins appear to be less structured or poorly folded.

**Figure 4.**
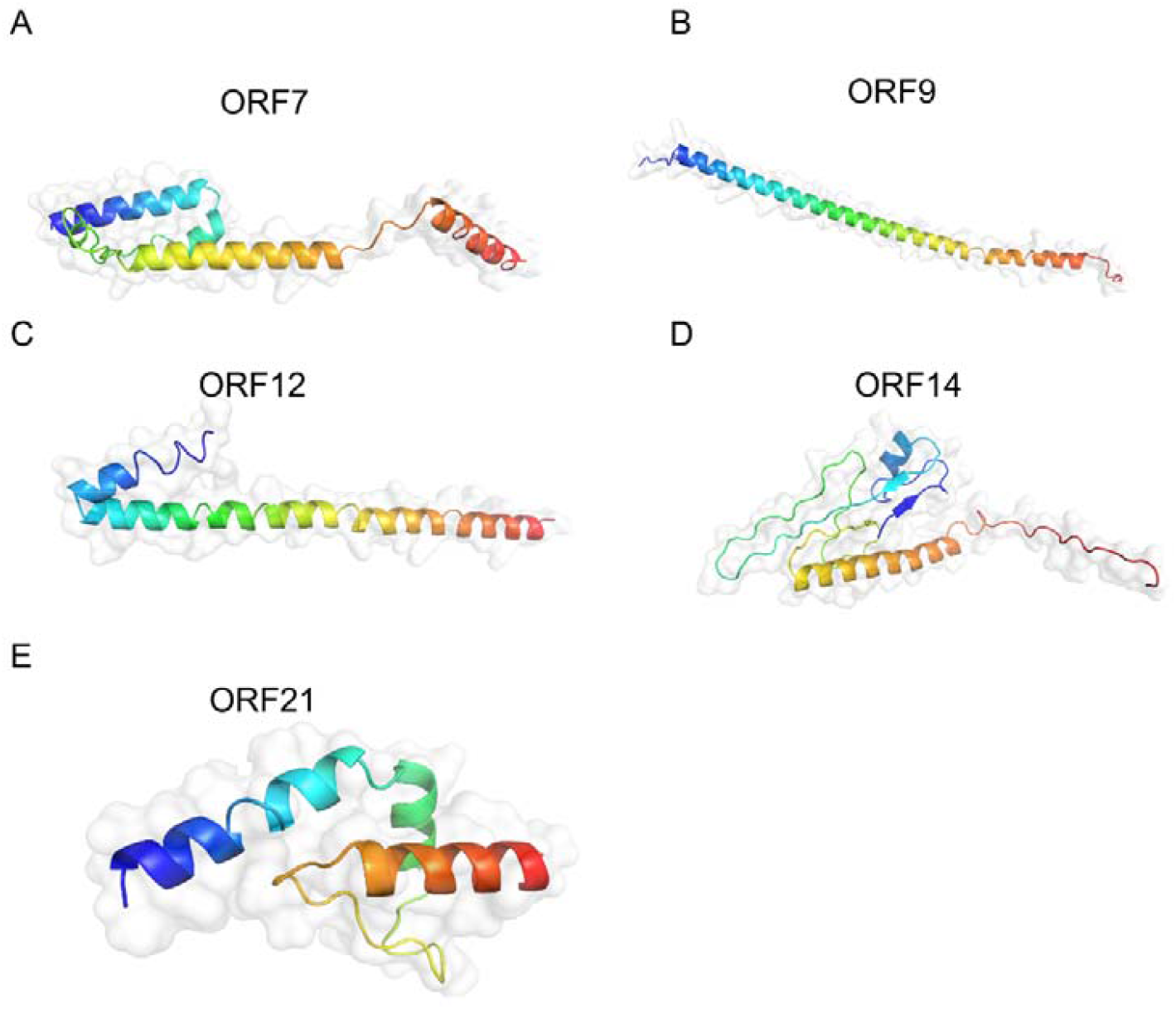
Structure prediction of nORFs using trRosetta. The structures were predicted in trRosetta using default parameters and were visualised in PyMol. The N-terminal of the nORF is present in blue colour and the C-terminal in red colour.

**Table 2** presents the properties of the five nORFs predicted to be translated based on kozak signals. HMI-PRED, HHpred, and motif search revealed the functional motifs in the proteins. While HMI-PRED showed the host proteins that could potentially interact with the viral protein. We also used DeepFold to identify the folds of the proteins, and we found that these novel proteins exhibit signalling and transcriptional folds.

**Table 2.** The physical and chemical parameters of nORFs were predicted using ProtPram (https://web.expasy.org/protparam/).

| S.No. | nORF | Length/Molecular weight | Theoretical pI | Aliphatic Index | GRAVY |
| --- | --- | --- | --- | --- | --- |
| 1 | ORF7 | 105 / 11616.41 | 9.44 | 80.86 | -0.164 |
| 2 | ORF9 | 102 / 11140.91 | 11.47 | 93.63 | -0.198 |
| 3 | ORF12 | 130 / 14472.74 | 9.56 | 89.15 | -0.122 |
| 4 | ORF14 | 79 / 8974.4 | 9.45 | 75.32 | -0.051 |
| 5 | ORF21 | 61 / 6928.04 | 5.99 | 107.5 | 0.67 |

**Supplementary File 4**, top 10 HMI-PRED hits for 5 nORFs showed that the proteins are involved in immune signalling, ion homeostasis, cytoskeletal organisation, signal transduction, apoptosis, immune regulation, and metabolism.

## Discussion

The genome is the blueprint of the functional capability and evolutionary history of a replicating entity. The composition and size of a genome provide important insights into the nature of a virus [26]. Obligate parasites, including many viruses and intracellular pathogens, tend to maintain compact genomes containing only the genetic material essential for survival and interaction with the host [27]. This genome reduction is shaped by evolutionary pressures to minimise replication time, reduce metabolic burden, and avoid detection by the host immune system [28]. Larger genomes not only require more resources and time to replicate but also increase the risk of mutations and errors in replication fidelity [29]. Although viruses have relatively small genomes compared to other replicating organisms, they utilise them with remarkable efficiency. Recent discoveries have revealed that many viruses possess a far more extensive coding repertoire than previously thought [30–32].

In this study, we wanted to explore the protein-coding potential of the RABV using an *in-silico* and multi-omics approach, and we report that there are functionally important ORFs on the antigenome. RABV synthesises a complete antigenome during an intermediate stage in the lifecycle of the RABV [10]. It is already reported that in RABV, a gene inserted in reverse orientation in the canonical strand can be expressed [10]. However, whether the RABV genome possesses any ORF in the reverse orientation (antigenome) is not known to our best knowledge. Here, we report that RABV has 16 nORFs on the antigenome, which can code for proteins longer than 50 amino acids; the smallest protein in our dataset was 51 amino acids in length, while the largest one encoded a 160-amino-acid protein. We used trRosetta to generate the structures of the nORFs, and we found that all of them folded into discernible higher-order structures. Interestingly, our analysis indicated that the RABV matrix protein, typically annotated as 202 amino acids in length, was predicted to have an additional 10 amino acids (MKKTGNTTDK) at its N-terminus. We sought to investigate if this extended form had been previously documented. Notably, several entries were found containing the N-terminal extension, and all these sequences (accessions OQ603609-OQ603685) originated from a recent study conducted in Georgia [33]. This observation raises broader implications regarding annotation consistency and the potential evolutionary and functional relevance of this extended protein isoform.

Furthermore, conservation of a coding frame is believed to hint towards the functional importance of that ORF [24]. We identify 5 nORFs (nORF7, nORF9, nORF12, nORF14, nORF21) or their parts that are conserved across the RABV clades (**Figure 2B**), which have diverged over a large evolutionary scale. The presence of the ORFs or domains across such large evolutionary scales suggests that the domains coded by these nORFs could have an important functional role associated with them. Recent independent studies have uncovered novel ORFs, often referred to as the dark proteome or orphan genes, in a variety of viruses and eukaryotes [34–36]. This suggests that such hidden coding regions in viral genomes may be more widespread than previously assumed.

To explore the functions of these nORFs, we used various in silico approaches. Using HHpred and Motifsearch tool (https://www.genome.jp/tools/motif/), we identified the domains present in the RABV nORFs, which were generally related to signalling as well as proteins involved in replication and transcription machinery. We also used DeepFold to generate the structures and then identified homologous proteins based on these structures (**Supplementary File 5**). We finally used HMI-Pred to find the putative interacting partners for these proteins. The 5 nORFs display domains for modulating diverse host cellular processes. Several nORFs show strong interactions with calmodulin (CALM1/CALM2), implicating them in calcium-mediated signalling, which can influence apoptosis, transcription, and membrane trafficking. Predicted mimicry of structural proteins like vinculin and plectin hints at interference with cytoskeletal organisation and cell adhesion, possibly to aid viral entry or spread. Collectively, these nORFs are likely to fine-tune host environments to favour viral replication and immune evasion through multifaceted molecular mimicry. However, elucidating the functional roles of these proteins requires further investigation into this largely uncharted coding space to better understand the organism’s biology.

In our proteomics data [15], we identified two nORFs supported by more than two unique peptides at high confidence, indicating that these nORFs are likely translated into proteins. Nevertheless, further proteomic analyses under varying conditions are needed to validate and understand the functional importance of these small proteins. Additionally, to investigate whether the nORFs are transcribed within host cells, we utilised transcriptomic data based on a strain (Tha-EU293111) to validate the expression of the nORFs [37] (data not presented). In this strain, we identified an ORF located within an intergenic region. Since the transcriptome data were generated using paired-end sequencing, capturing reads from both strands, this overlap made it difficult to confidently assess the expression of reverse-oriented nORFs in genic regions.

In conclusion, we report the identification of novel open reading frames in RABV, for which mRNA reads were detected, and peptides corresponding to these proteins were observed in the mass spectrometry data. We additionally validate the conservation of several ORFs across a long evolutionary scale. These observations suggest that RABV has more coding potential than previously reported, and these nORFs may play a crucial role in the virus’s pathogenesis. Additionally, the presence of nORFs is not unique to RABV, as other lyssaviruses also harbour more ORFs, and this phenomenon may also be common for other viruses.

## Acknowledgements

The authors thank Utpal Tatu lab members for their discussions and input.

## Author contributions

Conceptualisation: Ankeet Kumar and Utpal Tatu. Data curation: Ankeet Kumar.

Formal analysis: Ankeet Kumar. Investigation: Ankeet Kumar and Utpal Tatu. Methodology: Ankeet Kumar and Utpal Tatu. Supervision: Utpal Tatu.

Project administration: Utpal Tatu Resources: Utpal Tatu

Validation: Ankeet Kumar and Utpal Tatu. Writing – original draft: Ankeet Kumar.

Writing – review & editing: Ankeet Kumar, Utpal Tatu.

## Funding

Utpal Tatu acknowledges the DBT-IISc partnership. Ankeet Kumar acknowledges CSIR for financial support.

No external funding was obtained to support the project.

## Data Availability

The data supporting the findings in the study are available in the manuscript and as supporting data.

## Conflict of interest

The authors declare that they have no conflict of interest.

## Ethics approval

Not applicable

## Notes

### Competing Interest Statement

The authors have declared no competing interest.

