## Supplementary material for "Identification of Novel Open Reading Frames on the Rabies Virus Antigenome Strand and Assessing Their Expression": Supplementary_Data.docx

**Supplementary Table 1. The nORFs shared across two or more clades of RABV and their corresponding protein sequences.** The names of the nORFs were not consistent across the clades; therefore, the names were changed for ORFs for visualisation.

| **Cosmo** | **Africa-2** | **Africa-3** | **Arctic** | **Asian** | **Bats** | **Indian-Sub** | **RAC-SK** | **Notes** |
| --- | --- | --- | --- | --- | --- | --- | --- | --- |
| >ORF7 | >ORF7 | >ORF7 | >ORF7 | >ORF7 | >ORF7 | >ORF7 | >ORF7 |  |
| MLTCFWTDWKYFQVAKFLKSDGFSQIESKSITLETISFPPLMIAEGGSGCVPEAIRSFTSKRLLNTSLASGNMLRTALDIPPDPSPTTRQRDGKTLRSSRIGLSL | MFTCFWTDWKYFHVAKFLRSDGFSQIESRSITRETMSSPPLMIAEGGSGCVPEAIRSFTSNRLLNTSLASGNMLSTALDIPPDPSPTTRQRDGKTLRSSRIGLSL | MFTCFWTDWKYLQVAKFLKSDGFSHIESKSITLETMSSPPLIIAEGGSGCVPEAIKSFTSNKLLNTSLASGNMLSTARDIPPDPSPTTRHRDGKTLRSSRIGLSL | MFTCFWTDWKYLQVAKFLRSDGFSQIESKSITLETMSSPPLTIAEGGSGCVPEAIKSFTSNKLLNTSLTSGNMWSTALDIPPDPSPTTRHRDGKTLRSSRIGLSL | MFTCFWTDWKYFQVVKFLRSDGFSQIESKSTTLEIMSSPPLMIADGGNGCVPDAIRSFTSNKLLNTSLASGNMLSTALDIPPDPSPTTRHRDGKTLRSSRMGLSL | MESKSITLEMMSSPPLMIAEGGNGCVPDAIRSFTSNKLLNTSLASGNMLSTALDIPPDPSPTTRHKDGNTLSPSRIGLSL | MFTCFWTDWKYFHVVKFLRSDGFSQIELKSITRETMSSPPLMIAEGGSGCVPDAIKSFTSSRLLKTSLASGNMLSTALDIPPDPSPTTRHRDGKTLRSSRIGLSL | MESKSITLDMISSPPLMIAEGGNGCVPEAIKSFTSNRLLNTSLASGNILSTALEIPPDPSPTTRHKDGNTLRSSRIGLSL | ORF named as per Cosmo clade |
| >ORF10 | >ORF10 | >ORF10 |  | >ORF10 | >ORF10 | >ORF10 | >ORF10 |  |
| MTKYNLRFHDIRAKNRPSIFNSRSFGLRPIIKSSSGNPPRSIDFRNSRGLTGGLDRAVIITFSLGTGPPRFSDSQEASLVLVKECDLSSNISDGSIDSGISKICVIGSLCHVSPTKSTICLGGHVWVLI | MTKYSLRFHDIKAKNRPSILSSRSFGLRPIIKSSSGSPPRSIDFKNSLGLTGGLERAVMITFSLGTGPPLFSDSQEASLVLVKECDLSSSISDGSIDSGISKI | MTKYNLRFHDIKAKNLPSIFSSLSFGLRPIIKSSSGNPPRSIDFKNSRGLTGGLDRAVIITFSLGTGPPLFSDSQEANLVLVKECDLSSSISDGSIDSGISKICVIGSLCQVSPTKSTICLGGQV |  | MMTFSLGTGPPLFSDNQDASLVLVKECDLSSSISDGSIDSGISKICVIGSLCHVSPTKSTICLGGHV | MTKYNLRFQDIKAKNRPSIFSSLSLGLRPIIKSSSGSPPRSIDFRNSLGLTGGLERAVIMTFSLGTGPPLFSDSHDASLVLVKECDLSSSISDGSIDSGISKIWVIGSLCQVSPTKSTICLGGHVWVLI | MTKYNLKFQDIKAKNRPSIFSSLSFGLRPIIKSSSGNPPRSIDFRNSLGLTGGLDRAVIMTFSLGTGPPLFSDSQEASLVLVKECDLSSNISDGSMDSGISNICVIGSLCHVSPTRSTMCLGGHV | MIKSSSGSPPRSIDFKNSLGLTGGLERAVIMTFSLGTGPPLFSESHDASLVLVNECDLSSKISDGSIDSGISKIWVIGSLCQVSPTKSTICLGGHVWVLI | ORF named as per Cosmo clade |
| >ORF20 |  | >ORF20 | >ORF20 |  | >ORF20 |  |  |  |
| MYVNRAPGTKLVWYRVNREFPKTQWKNQKGYKQESLRNHLSLSLLRDVNSFFHIQEDQNVKVINLTLMKC |  | MMYVNRTPGTKFIWYRVNGEFPKAQRKTQKGYKQESLREHLSLSLWRDVNSFFHIQEAQYVKVINSTMIKCSDLTAP | MMYVNRAPGTKFVWYCVDGEFPETHWKSQKGHKQKSLRNHHSLSLLRDVDSFFHIQETPNVKVISLTSIKCSDLMAPLHMVFCSIAPKGWSQARMFSPPEVNKLETCGI |  | MMYVNGTPGAELVWYRVDGESPETQRGDQERHIQKSLRDHLPLGPLRDVNSFFHI |  |  | ORF named as per Cosmo clade |
| >ORF13 |  | >ORF13 | >ORF13 |  |  |  |  |  |
| MSLIVFDISIDQINSFFSHLKIGQTGSNSTRIDKDLAHVWGGSKGGVLVFFMMDIHNP |  | MSLIVFYIPVDQINGLFSHLKISQTGSNCTRIDKDFAHIGDGSEGGVLVFFMRDIHSR | MSLIVFYVFVDQINSLLSHLQISQTGSNCTRIDKDFAHIWGGSKGGVLVFFMKDIHNLWISCVPLFNYLRVTRICLV |  |  |  |  | ORF named as per Cosmo clade |
|  | >ORF14 | >ORF14 |  | >ORF14 |  |  |  |  |
|  | MCWLGTCQMPYRSQPFQGNSIPPPGGFSALYSQVYNLAASLWGKTWNRNGSGQLCISCSC | MYWLNTSQTPCHSRPSQGSSTLPLGGFLALYFQVYNSTASLWGKIWNRNGLDRLYIFCFCSTPSGHHITRERSLTVQQFRPL |  | MYWQDTSRMPYHSQLFQGNSTPLLGEFLVLYFQVCNLITSLWVKTLNRNGPSQPCTSDSCLTQSGHHITRERSSTGRQFHLL |  |  |  | ORF named as per Africa-2 |
|  | >ORF11 | >ORF11 | >ORF11 |  |  |  |  |  |
|  | MFNPGEIHVVCSGQESEVRYSTPVSAHFMSCHQSMMFHNFDERGCLKNLLYPICNVCLVVACVLSRYFA | MLNPGEKHVVGPSQESEVWYSTPISTHLMSCHQGMMFHDLDERGRLENLLYPICNVCLIIAGVLA | MLNPGEKHVVGSCQKSEVWYSAPVGTHLVSCHQSMMFHNLNKRGCLENLLYPICNVCLIVTGVLPRYFA |  |  |  |  | ORF named as per Africa-2 |
|  | >ORF6 |  | >ORF6 | >ORF6 |  |  |  |  |
|  | MFFLPVSSFSGTYSGGGSHRSSSSGGADTGEGFWVSSSLQFFTILRMKFILSVVLPVFFMLNLEQLRFNCISGLSLKVWSAGCITI |  | MQKFLRFFLPVSSFSGTYSGGGSHRSSSSGGADTREGFWVSSSLQFFTILRRKFILSVVLPVFFMSNSERFIFYRLSGPSIKA | MFFLPVSSFNGIYSGGGSHRSSSSGGADTGEGFWVSSSLQFFTILRRKFILSVVLPVFFMSGPEQLRLYRLTELSLKICQQDV |  |  |  | 17, 11, and 10 were already taken. So a new ORF name was given to all |
| >ORF9 |  |  | >ORF9 |  | >ORF9 |  |  |  |
| MIDSGIPKKDELNSSLRNRGNRGSTDVKKSIKLSLWVLDNRIASRNSEFSTLSTSSYKAFLIASLSRMVGLAPPLIFKVVGSSRRRVKLSSVLSPRSGFPAS |  |  | MMDSGIPRKDELKSSLRNRGNKGSTDFKKSIKLSLWVLDNKIASLNSEFSTLSTSSYRAFLIASLSKMVGLAPPLIFKAVGSSRRRVKLSSVLSPRSGFPAS |  | MIDSGIPKKDELKSSLRNRGNKGSTDFKKSIKLSLWVLDKRIASLNSEFSTLSTSSYRAFLIASLRRMVGLAPPLIFRVVGSSRRRVKLSRVLSPRSGFPAS |  |  | ORF named as per Cosmo clade |
| >ORF18 |  | >ORF18 |  |  |  |  |  |  |
| MPVTAIHKNQVMYVVELIQSPKEPSCTGTERLFELTYLVFYKYRKVSQGMNERPKLFCSLD |  | MPVTAINKNQVMDVVELIQNLKEPFCTRTESPFKLTHLIFYEYRKVSQGVNERPKLLCPLD |  |  |  |  |  | ORF named as per Cosmo clade |
|  | >ORF15 |  |  | >ORF15 |  |  |  |  |
|  | MIIGVWIVQRLFISRVTGHLPVVCSPTRVRCRAKVLSPERGGDITDEVSISFCFCHHACTSKPVHFYG |  |  | MIIGIRIMERFLISGVTCHLPVICCSACIWCRAEVLSLKRGGDIANKVCVGLCFCHHTRASEPVHFYG |  |  |  | ORF named as per Africa-2 |
| >ORF12 | >ORF12 |  | >ORF12 | >ORF12 |  |  |  |  |
| MTLESTQLHGVQGGLLNPGFFSKRTPQGEVIFRTWIVERTASSHSPVSPPLLYDSHEDMIFPLWGVTDTSLPVPLRLCCVGSDRLTLLQHVIRKIINIKAVRAPALSNTYFPQFGRPKSTPEIRSCTSGR | MTGFHSLVSPPLLYDSHEDITFPFWEVTDTFLPVPLILCCVDSGRLIFLQHVIRKIISIRAIRAPALIKTYFPQVGRPRSTPEICLCTSGRCTSTKSSASSSSLKTVDGSAKGCIRGITEDSNNSICC |  | MTGLHSLVSPPLLYDSHEDMTFPLWEVADTFLPIPPRLCCMDLGRLPLPQHVIKKTVNIMAIRTPAIIKTYFPQLGRPRSTPEICLCTSGR | MACFYSLVSPPLLYDSQEDMTLPLWEVTDTFLPVSPRRCCIDSTPLAFLQHVIKKIINISAIRAPTLTSTYFPQFGRPRSTPETCLWTSGR |  |  |  | ORF named as per Cosmo clade |
|  |  | >ORF16 |  |  |  | >ORF16 |  |  |
|  |  | MTLSRTHSYLQSDKFCVVIPGDLLRVLQLLPLQRAPGVSPLEDEGSPQFVYPVHALRDWSS |  |  |  | MALSRTNPYLQSDKLRVIIPGDLLRVFQFLPFKRASRVSPLEDESSPQFIYPIHALWDWSS |  | ORF20 was taken, so ORF16 was given to this ORF |
| >ORF17 | >ORF17 |  | >ORF17 | >ORF17 |  | >ORF17 |  |  |
| MILLSLLDSTNNWNFLELAKATHPSTHLRGSLDPSRAKRVTPGTFFASLTISSRFIFNCSKLYKSIPEDREGNLYFLEKLSAIW | MTLFSLSGSTNNWNLLELARATHPSTHLRGSLDPSWAKRVTPGTFFASLTMSSRFIFNCSKLYKNIPDDLEGNLYFLEKLSAIW |  | MILFSLSGSTNNWNFLELARPTHPSTHLRGSLDPSWARRVTPGTFFASLTMSSRFIFNCSKLYRNIPDDRDGNLYFLEKLSATW | MILFSFSGSTNSWNFLEFARATHPSTHLRGSLDPSRAKRVTPGTFFDSLTISSRFIFNCSKLYKNIPDDRDGNLYFLEKLSORFIW |  | MILFSLSGSTNNWNFLELARATHPSTHLKGSLDPSWARRVTPGTFFDSLTMSSRFIFNCSKLYKNIPDDRDGNLYFLEKLSAIW |  | ORF named as per Cosmo clade |
|  |  |  | >ORF21 |  |  |  | >ORF21 |  |
|  |  |  | MLNTLLEKTIIMAIMIATLDKVPLRNSRSSTILWTKKDSTSLSIRVMIISL |  |  |  | MVLQQILKCLKILGSDGGLNKGSVNGFETLNTLLEKTIMMATIMDTLDKVPRCNSRSSTIL | ORF named as per Arctic clade |

**Supplementary Table 2. The Kozak sequences identified for nORFs using ATGpr tool.**

| Novel ORF | Identity to Kozak rule A/GXXATGG | Length | Sequence |
| --- | --- | --- | --- |
| nORF12 | cXXATGa | 130 | MTLESTQLHGVQGGLLNPGFFSKRTPQGEVIFRTWIVERTASSHSPVSPPLLYDSHEDMIFPLWGVTDTSLPVPLRLCCVGSDRLTLLQHVIRKIINIKAVRAPALSNTYFPQFGRPKSTPEIRSCTSGR |
| nORF7 | GXXATGt | 105 | MLTCFWTDWKYFQVAKFLKSDGFSQIESKSITLETISFPPLMIAEGGSGCVPEAIRSFTSKRLLNTSLASGNMLRTALDIPPDPSPTTRQRDGKTLRSSRIGLSL |
| nORF9 | cXXATGa | 102 | MIDSGIPKKDELNSSLRNRGNRGSTDVKKSIKLSLWVLDNRIASRNSEFSTLSTSSYKAFLIASLSRMVGLAPPLIFKVVGSSRRRVKLSSVLSPRSGFPAS |
| nORF14 | cXXATGc | 79 | MLRDSGVSCQFHASCQSPIPFYISTFYLHQRTWGDLISFSCNHDSIAGPVFRTCPLKKLHCRRQIGTYIVRIKFGGAHA |
| nORF21 | GXXATGa | 61 | MTLFCTHSYLQSDELSIIIPSDLLRVFQLLPFKRAPGISPLEDKGSPQLVYPVHALRDWSS |
